# Protection of the prodomain α1-helix correlates with latency in the transforming growth factor-β family

**DOI:** 10.1101/2021.06.24.449816

**Authors:** Viet Q. Le, Bo Zhao, Roxana E. Iacob, Yuan Tian, Cameron Toohey, John R. Engen, Timothy A. Springer

## Abstract

The 33 members of the transforming growth factor beta (TGF-β) family are fundamentally important for organismal development and homeostasis. Family members are synthesized and secreted as pro-complexes of prodomains that are non-covalently bound to the growth factor (GF). The pro-complexes of some members are latent and require activation steps to release the GF for signaling. Why some members are latent while others are non-latent is incompletely understood, but crystal structures and hydrogen-deuterium exchange (HDX) of four family members have begun to unravel how latency is regulated. Here, we extend this understanding by comparing pro-complex conformation in negative stain EM (nsEM) and HDX of ActA, BMP7, BMP9, BMP10, GDF8, TGF-β1, and TGF-β2. nsEM revealed that family members varied in either adopting cross-armed, open-armed, or V-armed configurations. Latency was achieved in both cross-armed and V-armed but not open-armed conformations. HDX revealed remarkably varying patterns of exchange between family members, consistent with large prodomain sequence divergence. We observed a strong correlation between latency and protection of the prodomain α1-helix from exchange, which in latent members coincided with greater buried surface area of the α1-helix and more hydrogen and cation-pi bonds from the prodomain fastener and GF to the α1-helix. Strong sequence conservation of the α1-helix and fastener only in latent members suggests that similar interactions are conserved and sufficient to confer latency. Moreover, most members exhibited rapid exchange in the unstructured “association region” at the prodomain N-terminus, highlighting their availability for interacting with factors that may regulate latency and extracellular storage.

## INTRODUCTION

The TGF-β family comprises 33 genes which encode TGF-βs, bone morphogenetic proteins (BMPs), growth and differentiation factors (GDFs), and activins/inhibins. Family members regulate homeostasis, establish the anterior-posterior, ventral-dorsal, and left-right axes during embryonic development, and direct fine details of skeleton and organ development (1,2). In general, members are synthesized as a proprotein composed of a large, divergent N-terminal prodomain and a smaller, more conserved C-terminal growth factor (GF) domain, separated by a proprotein convertase (PC)/furin cleavage site. Prodomains are required for growth factor domain folding and dimerization (3–5), storage in the extracellular matrix (ECM) or on cell surfaces (6–17), and regulate growth factor activity and signaling range (17–24).

Subsequent to disulfide bond formation and dimerization in the ER, family members are cleaved by PC/furin, usually in the Golgi, and secreted as pro-complexes of noncovalently associated prodomains and growth factor dimers. Pro-complexes of most members are active, i.e., the prodomains can be displaced by receptor binding. In contrast, isolated pro-complexes of a few family members such as TGF-βs 1–3, GDF8 (myostatin), GDF9 in some species and not others (25), and GDF11 are latent (25–34). Activation of latent TGF-βs 1 and 3 is mediated by αV integrins that bind to an RGD-motif in the prodomain and transmit actin cytoskeletal force that distorts the prodomain and liberates the growth factor (35–37). Activation of GDF8 and GDF11 is mediated by cleavage of the prodomain by Tolloid (TLD) metalloproteases (33,38). The mechanisms governing latency and non-latency are not fully understood. Although BMPs are not latent in vitro, mutational and genetic studies show that the distance over which BMP signaling spreads in vivo is regulated by “shadow” PC cleavage sites in the prodomain (18–20,24). Furthermore, BMPs are stored in tissues in vivo (8,9,39–43), suggesting that the growth factor is not released immediately after secretion. Moreover, interaction with the ECM component fibrillin has been reported to confer latency on the BMP7 pro-complex (44). Better understanding of the properties of GF-prodomain complexes is therefore essential to understand the biology of GF signaling.

The overall factors that regulate latency in the TGF-β family are incompletely understood. Our previous structural and conformational dynamics studies of GDF8 activation provided a framework for understanding its latency (45). TLD cleavage of latent GDF8 yielded prodomain fragments that remained associated with the GDF8 growth factor in a complex “primed” for dissociation (i.e., primed GDF8). Hydrogen-deuterium exchange coupled to mass spectrometry (HDX-MS) showed that TLD cleavage increased deuterium exchange in a small subset of peptides that map to prodomain–GF interfaces (45,46) and thus defined regions with a role in maintaining latency. Given that similar prodomain–GF interfaces are present in structures of latent TGF-β1 and non-latent BMP9 and ActA (23,47–49) and are expected to be conserved in other TGF-β family members, the relative strength of these interfaces may be an important factor underlying latency. Here, we characterize a diverse set of latent and non-latent TGF-β family members using negative stain electron microscopy (nsEM) and HDX-MS to determine how overall pro-complex conformation and the structural stability of prodomain-GF interfaces relate to latency and non-latency.

## RESULTS

### Negative stain EM shows three overall pro-complex conformations

We used negative stain EM (nsEM) to characterize the overall conformations of latent TGF-β2 and non-latent BMP10 pro-complex dimers and compared them to previously published EM structures of related family member dimers. TGF-β2 adopted a ring-like conformation (termed cross-armed) similar to TGF-β1 (47) (Fig. 1A,C). In contrast, BMP10 predominately adopted a nearly linear open conformation (termed open-armed) similar to its closest homolog, BMP9, and was also similar to BMP7 (23) (Fig. 1B,D,&F). In comparison, previous nsEM of latent GDF8 revealed a V-shaped conformation (termed V-armed) that is intermediate in arm domain orientation between the cross-armed conformation of the two TGF-βs and the open-armed conformation of the three BMPs (45) (Fig. 1E).

**Figure 1.**
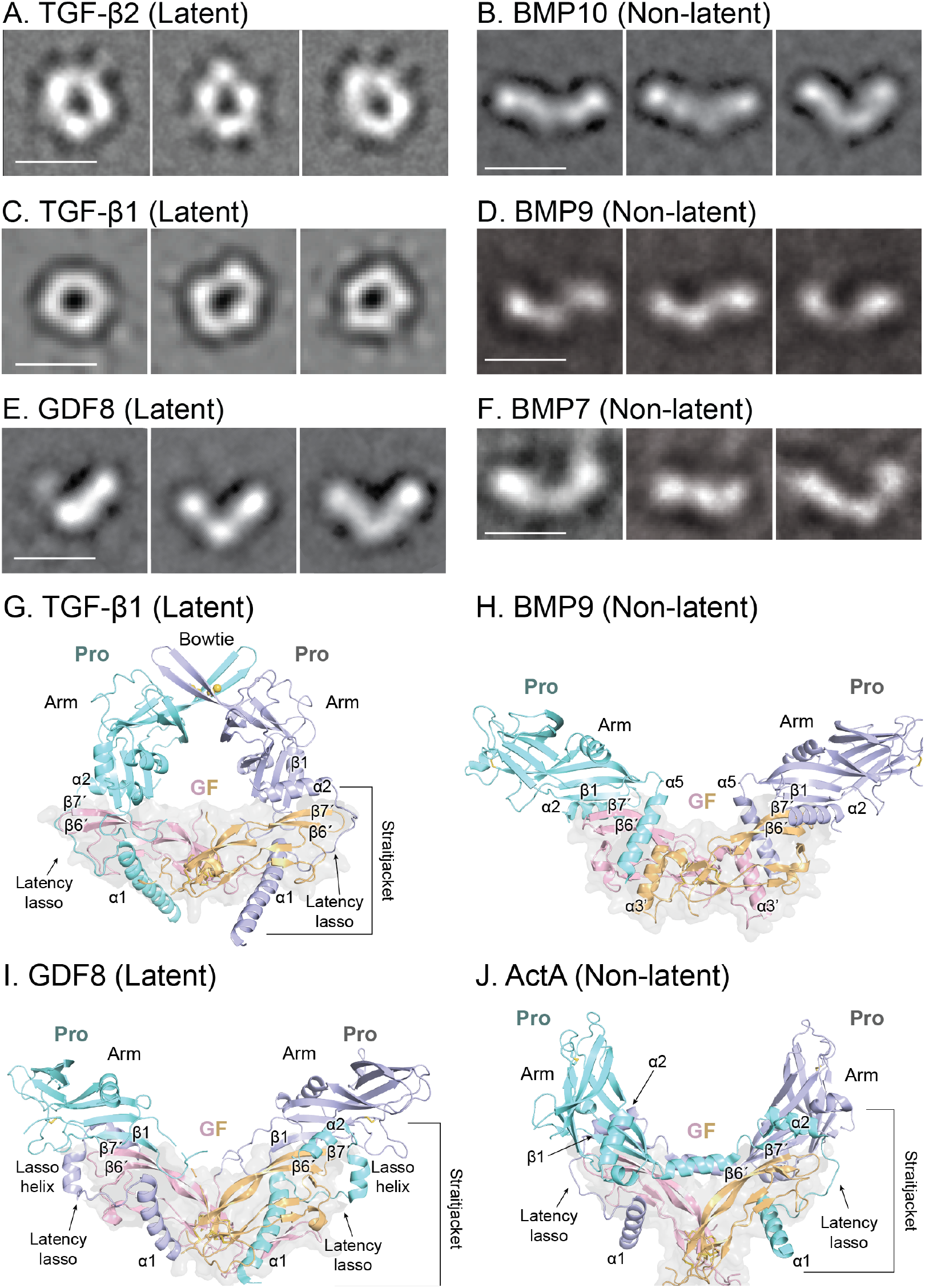
Structure and conformation of TGF-β family member pro-complexes. **A–F.** Representative negative stain EM class averages of TGF-β2 and BMP10 compared to previously published class averages of TGF-β1, BMP9, GDF8, and BMP7 (23,45,47). Scale bars = 10 nm. **G–J.** Crystal structures of TGF-β1, BMP9, GDF8, and ActA superimposed on their GF dimers (23,46,48,49). Yellow spheres indicate the position of the bowtie disulfide bonds in TGF-β1 (G). The solvent accessible surface of the GF is shown as a transparent gray surface. Pro = prodomain. GF = growth factor.

The conformations of the six different TGF-β family member dimers in nsEM correspond well with those observed in representative crystal structures (23,46–49) (Fig. 1G–J). The cross-armed conformation of TGF-β2 and β1 matches the crystal structure of TGF-β1 (Fig 1A,C,&G), in which the arm domains of each prodomain monomer are disulfide linked to one another at their tips distal from the GF domains. On the opposite side of the ring from the arm and bowtie, the latency lasso and α1-helix wrap around the growth factor dimer to form the other half of the ring. The open-armed conformations of the three BMPs resemble the BMP9 structure, in which the arm domains are oriented with their tips pointing away from one another (Fig. 1B,D,F,&H). Meanwhile, the V-shaped particles of latent GDF8 in nsEM correlate well with the pro-complex crystal structure of latent GDF8 (46) (Fig. 1E&I). The crystal structure of BMP9 has a more open vee, i.e. is more open-armed, than GDF8, which in turn is more open than the vee of ActA (Fig. 1H-J). nsEM showed heterogeneity among BMP class averages of the vee angle (Fig. 1D, E, F) and some BMP9 class averages even showed an S-shape (Fig. 1D). This heterogeneity may reflect genuine conformational flexibility and also effects of adsorption onto the EM grid.

### TGF-β family pro-complexes exhibit disparate patterns of HDX overall

Next, we used HDX-MS to investigate the structural dynamics of TGF-β1, TGF-β2, BMP7, BMP9, BMP10, and ActA at pH 7.5. Previously published HDX data for GDF8 (45) at pH 7.5 are also reported for comparison. Overall, we obtained 73–90% sequence coverage (Table S1) with nearly complete coverage of the prodomains and less coverage of the growth factor domains (Fig. 2); the disulfide-rich growth factor domains are difficult to reduce in HDX quench conditions, which results in incomplete pepsin digestion. HDX results are displayed for aligned sequences at two different time points in Fig. 2 (results for all time points and all peptides are found in Figs S2-S4) and displayed on crystal structures of TGF-β1, GDF8, ActA, and BMP9 in Fig. 3. As nsEM showed that TGF-β2 had a structure similar to TGF-β1, and BMP7 and BMP10 had structures similar to BMP9, the alignments in Fig. 2 were used to display the HDX data on the corresponding crystal structures in Fig. 3. All comparisons of backbone amide HDX between TGF-β family proteins is qualitative because our HDX-MS measurements were of relative deuterium incorporation and were not corrected for back-exchange during analysis (maximally deuterated controls could not be prepared for these proteins). HDX of related proteins with divergent sequence can be compared [more details can be found in (50–52) provided the interpretation remains mostly qualitative, as has been successfully done previously for many systems including GDF8:TGF-β1 (45), integrins (53) heme-oxygenases (54), Ras-family proteins (52), processivity clamps (52), and allelic variants of HIV Nef (51).

**Figure 2.**
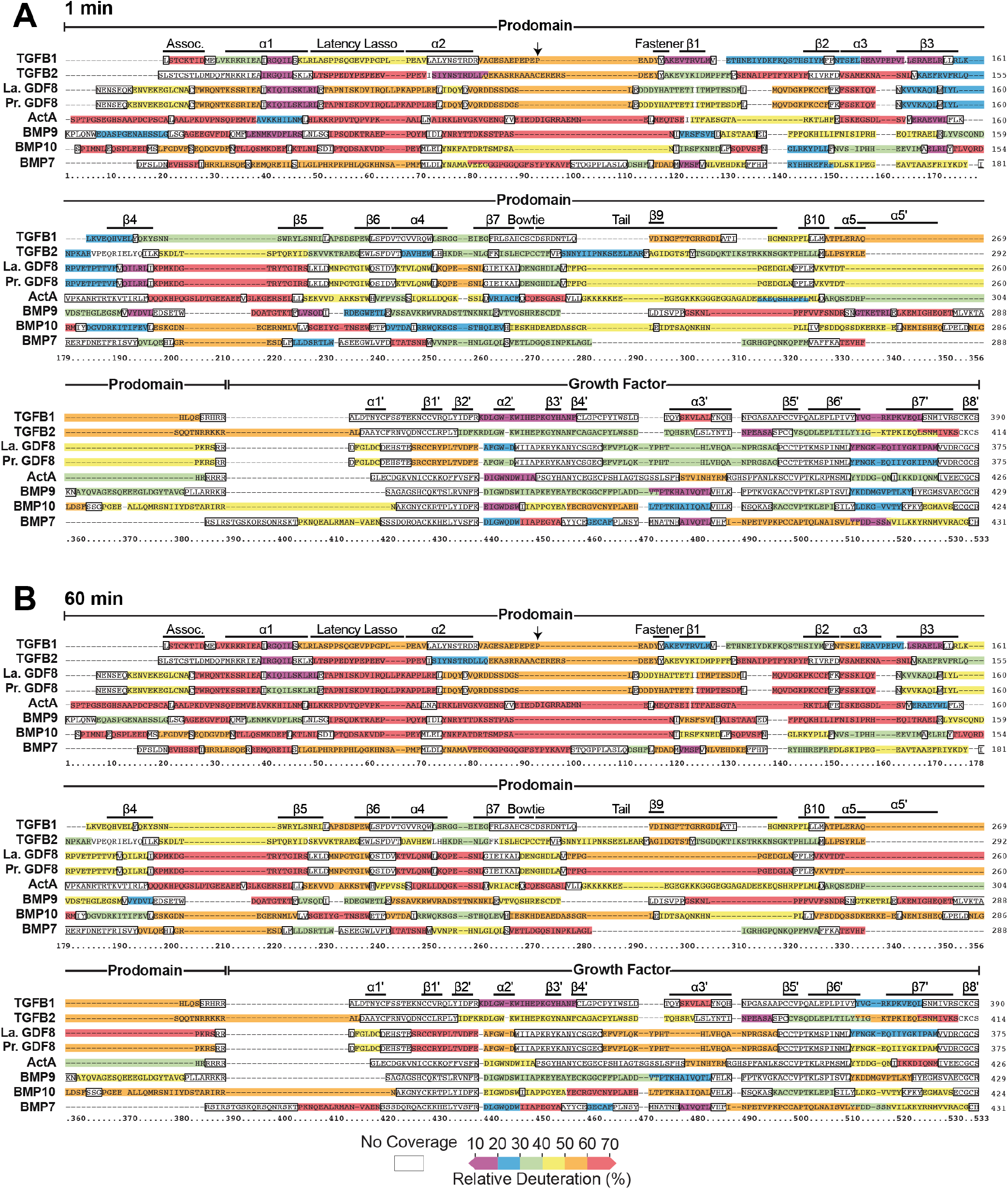
Hydrogen-deuterium exchange of TGF-β family members indicated on their sequences. **A and B.** Deuteration at 1 min (*A*) and 60 min (*B*). Peptides are colored according to the key shown. The data are overlaid onto a sequence alignment of all proteins investigated that has been corrected to align structurally homologous positions. Prodomain and growth factor boundaries and structural elements are marked above the alignment. Dashes represent gaps in the sequence alignment and are colored if sequences at each end are included in a peptide covered by HDX-MS. Regions that lack HDX peptide coverage or correspond to the rapidly back-exchanging N-terminal residue of peptides are shown as unfilled rectangles. A downward arrow indicates the position of the Cys residue found in TGF-β2 but not in TGF-β1.

**Figure 3.**
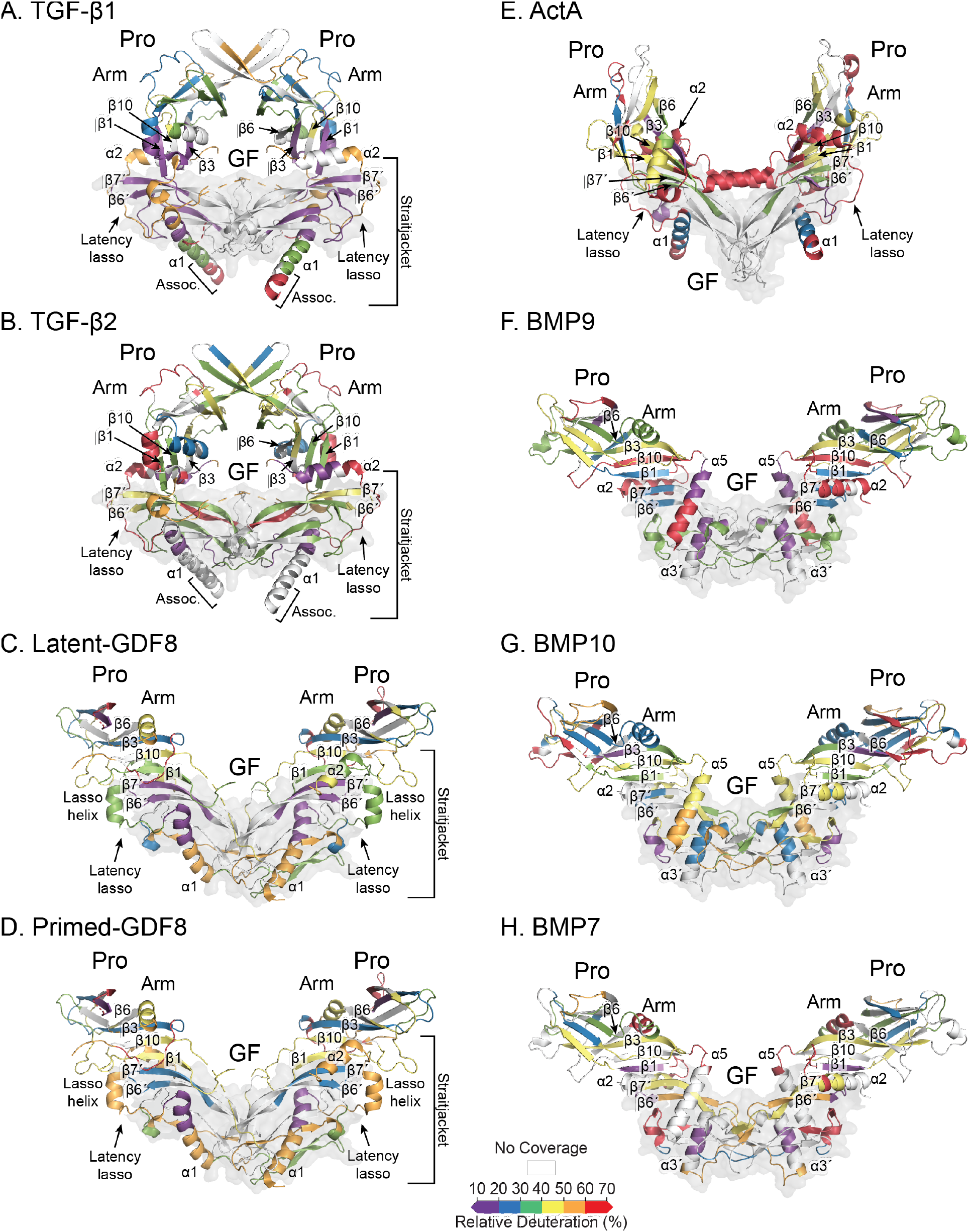
Hydrogen-deuterium exchange of TGF-β family members indicated on structures. **A-H.** Deuterium exchange at 1 min is colored onto ribbon diagrams of crystal structures or onto crystal structures of the closest homologue, according to the key and the alignments shown in Fig. 2. TGF-β2 utilizes the TGF-β1 structure and BMP7 and BMP10 utilize the BMP9 crystal structure. The GF is shown as a gray transparent surface as in Fig. 1.

Strikingly, overall exchange varied greatly even among the most closely related members studied here. Exchange was faster almost everywhere in TGF-β2 than β1, although HDX in the β9 to β10 region in TGF-β2 was slower than in TGF-β1 (Fig. 2). Differences were also pronounced among BMP7, 9, and 10. These differences were consistent with the large sequence variation among prodomains. For example, the TGF-β1 and β2 GFs are 71% identical while their prodomains are 39% identical; the BMP9 and BMP10 GFs are 64% identical while their prodomains are only 33% identical. As dissociation of the prodomain from the GF is required for Type 1 and Type 2 receptor binding, this remarkable variability in exchange suggests variation in regions of the prodomain that are most susceptible to breathing movements (fast exchange) with important implications for GF dissociation, i.e. activation.

Variability in exchange among family members was evident throughout their prodomains. The first portion of the prodomain is called the association region because in TGF-β1, β2, and β3 it contains the conserved Cys residue that disulfide links to a milieu molecule that mediates pro-complex storage in either the ECM or on cell surfaces (13,14,16,17,55). Furthermore, the complex crystal structure of TGF-β1 linked to the milieu molecule glycoprotein A repetitions predominant (GARP) shows that each TGF-β1 monomer forms a buried interface with GARP that involves varying lengths of amino acid residues flanking the conserved Cys, e.g., ^30-^LSTCKTID^−37^ in one monomer and ^32-^TCKTI^−36^ in the other (56). The longer 8-residue sequence was used to define the association region in Figs. 2 and 3. No milieu molecules were present in our study, and in the absence of a partner, the association region of TGF-β1 was rapidly deuterated (peptides corresponding to the association region of TGF-β2 were not covered in HDX). The N-terminal segment containing the association region is often longer in other family members and is unstructured in GDF8, ActA, and BMP9 crystal structures (23,46,49) which have no known milieu molecules that associate in this region. The N-terminal peptides in other family members exhibited moderate to rapid HDX, with the exception of BMP9, which exchanged much more slowly throughout the time course.

The α1-helix immediately following the association region also differed in exchange between family members. In TGF-β1, TGF-β2, and latent GDF8, peptides from the C-terminal half of the α1-helix were strongly protected from exchange. In contrast, similar α1-helix peptides from the activated form of GDF8 (primed GDF8), ActA, and the three BMPs exhibited moderate to fast exchange at the 60-minute time point (Fig. 2B, Fig S3-S4). In crystal structures of TGF-β1, ActA, and GDF8 pro-complexes, the α1-helix has a similar conformation (Fig. 3); the slower HDX of TGF-β1 and GDF8 than ActA are consistent with greater burial and polar noncovalent bonds as described in Discussion. In contrast to TGF-β1, ActA, and GDF8, in BMP9 pro-complex crystal structures no density is apparent for the α1-helix; the α5-helix appears in a similar location but with a distinct orientation.

The α1-helix is linked to the latency lasso, a rapidly exchanging loop in all family members studied here (Fig. 2), followed by the α2-helix to form the straitjacket element that encircles the growth factor domain in structures of TGF-β1, GDF8, and ActA (Fig. 3). The α2-helix forms a conserved interface with the convex surface of the growth factor domain on the side opposite to the cleft inhabited by the prodomain α1-helix (Fig. 3). Slow exchange of the α2-helix was only apparent in TGF-β2, although peptides corresponding to the α2-helix in TGF-β1 were not covered.

HDX differences extend into the bulk of the prodomain, which is composed of two conserved 4-stranded antiparallel β-sheets that form a jellyroll fold in pro-complex crystal structures and is referred to as the arm domain (Fig. 3) (23,46–49). The proximal β-sheet is composed of β-strands 1, 3, 6, and 10, whereas the distal β-sheet is composed of β-strands 2, 4, 5, and 7. The β1-strand initiates the arm domain and in TGF-β1 resides in a region that exchanged more slowly relative to the other family members. The β3-strand exhibited variability with slow exchange in TGF-β1 and ActA compared to moderately faster exchange in latent and primed forms of GDF8, BMP7, and BMP10 (Fig. 2B). Comparisons of peptides from the core of the β4-strand region showed slower exchange in BMP9 compared to latent and primed GDF8. The β5-strands of BMPs 7 and 9 were more protected from exchange than in BMP10 and ActA.

HDX differences continued into the growth factor domains, particularly the α3’ helix and the β6’-β7’ strand regions. Slow exchange of the α3’ helix was observed for BMP7 and BMP9, whereas the α3’ helices of TGF-βs 1 and 2, GDF8, and ActA experienced moderate-to-fast exchange. In pro-complex structures, the α3’ helix varies in structure and position. In BMP9, the α3’ helix is well formed, whereas in GDF8 and ActA it is disordered, correlating with the HDX results. The α3’ helix varies greatly in order or disorder among different TGF-β1 crystal structures and in orientation among different examples in crystal structures, suggesting it is dynamic in pro-complexes (37,47,48,56). The β6’ and β7’ strands of the growth factor domain form part of the concave surface to which the prodomain α1-helix of TGF-β1, ActA, and GDF8 or the α5-helix of BMP9 binds (Fig. 3,4). Slow exchange was observed in the C-terminal end of the β6’ strand and the N-terminal half of the β7’ strand of TGF-β1 and latent GDF8, whereas faster exchange was observed for this region in the remaining family members including, notably, TGF-β2 (Figs. 2&3).

**Figure 4.**
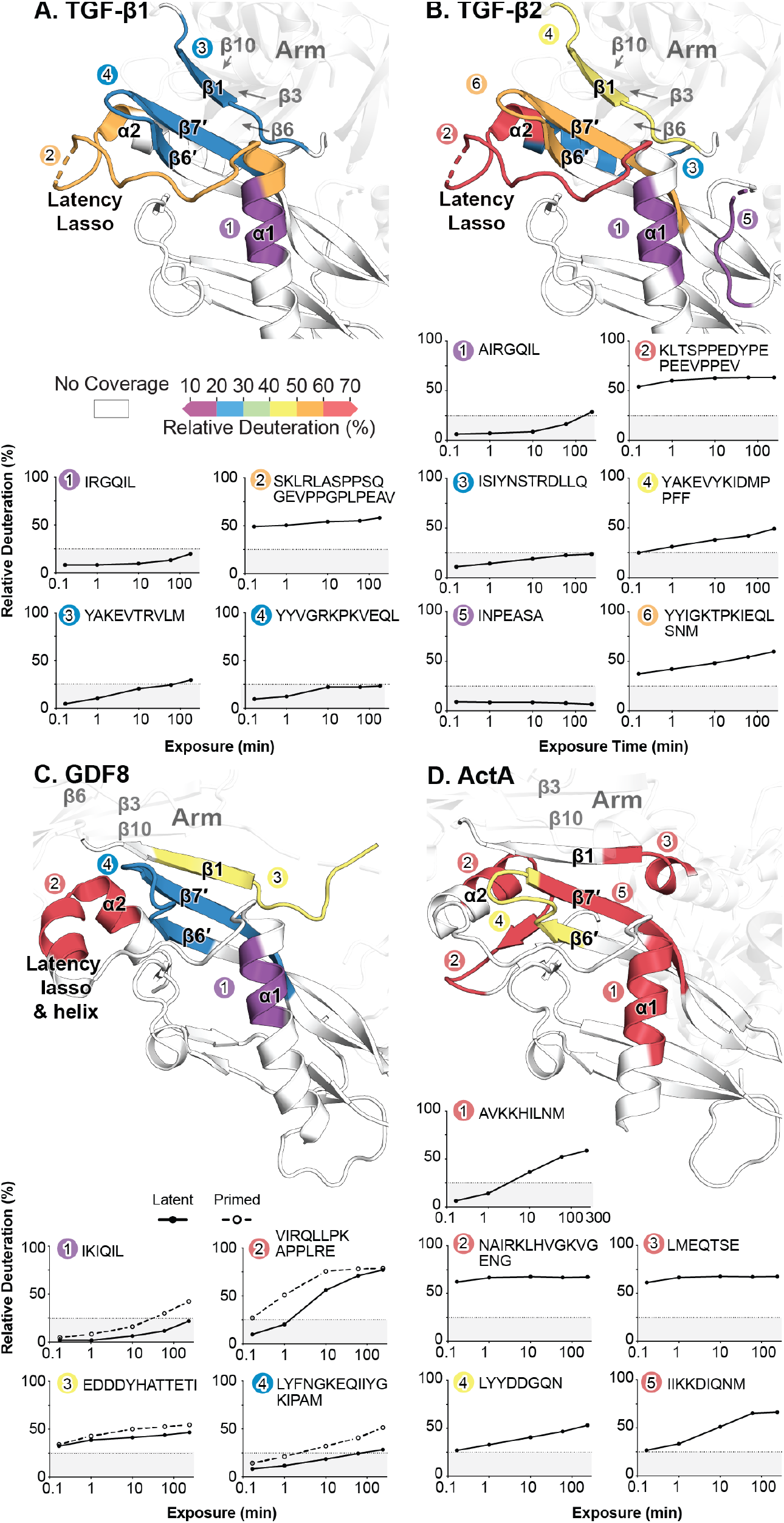
HDX in straitjacket–growth factor interfaces. **A-D.** Interfaces are shown for the indicated family members. In the top half of each panel, peptides are colored according to exchange after 1 h, using the key shown. In the bottom half of each panel, HDX uptake graphs are shown for the same peptides keyed by number and color. Gray boxes indicate percent deuteration in the 0–25% range. HDX data for latent and Tolloid-cleaved GDF8 are plotted as solid and dashed lines, respectively.

### Latency correlates with slow exchange in the C-terminal portion of the prodomain α1 helix

Despite substantial variation in HDX in other regions, a clear correlation was observed between slow exchange of the prodomain α1-helix and latency. The C-terminal half of the α1-helix in latent TGF-β1, TGF-β2, and GDF8 was highly protected from exchange (Fig. 2B, peptide 1 in Fig. 4A-C). However, this region of the α1-helix in non-latent Activin A, BMPs 7, 9, and 10, and primed GDF8 exchanged more rapidly (Fig 2B, peptide 1 in Fig. 4C,D). The GF β6′- and β7′-strands, which interact with this region of the α1-helix, also exchanged slowly in TGF-β1 and GDF8 (peptide 4 in Fig. 4A&C); however, TGF-β2 was an exception; it exchanged more rapidly than several non-latent family members (peptide 6 in Fig. 4B and Fig. 2).

## DISCUSSION

Our nsEM and HDX-MS studies presented here offer important insights into the diversity of pro-complex structures and mechanisms underlying latency in the TGF-β family. Extension of nsEM studies to TGF-β2 and BMP10 pro-complexes, which were previously uncharacterized structurally, revealed a cross-armed conformation for TGF-β2 and an open-armed conformation for BMP10. Together with our previous nsEM work, these results further highlight the existence of at least three pro-complex conformational states: cross-armed, open-armed, and V-armed. Cross-armed pro-complexes have thus far only been found to be latent, i.e., TGF-β1 and β2, whereas V-armed pro-complexes can be either latent (GDF8) or non-latent (ActA). Thus far, open-armed conformations have only been found to be non-latent as illustrated by BMP7, 9, and 10.

Importantly, our comparative HDX studies on multiple family members revealed that slow exchange in the C-terminal portion of the prodomain α1-helix correlated strongly with latency. Latent TGF-β1, TGF-β2, and GDF8 showed slow exchange in this region whereas non-latent ActA, BMP7, BMP9, and BMP10 all showed moderate to fast exchange at 60 minutes. The α1-helix is a key region of the prodomain that interacts with the GF, and slow exchange in both the prodomain α1-helix and GF β6′–β7′ region of TGF-β1 and GDF8 suggest that these elements form a stable interface that contributes to strong prodomain–growth factor binding. Meanwhile, the faster exchange of the GF β6′– β7′ region in TGF-β2 compared to TGF-β1, suggests a less stable interface with the α1-helix, and that perhaps other structural features of TGF-β2 compensate to maintain latency (discussed below). Taken together, our studies strongly suggest that a protected prodomain α1-helix occupying the GF hydrophobic cleft is a signature feature of latency.

Among latent members studied, TGF-β2 is not only distinguished by a faster exchanging GF β6′-β7′ region but also by faster dynamics overall. Compared to TGF-β1, TGF-β2 has a much longer prodomain with 283 residues compared to 249 in TGF-β1 and many insertions and deletions (Fig. 2). This larger size may contribute to latency. Several candidate regions emerge that may also contribute to TGF-β2 latency. The first is slow exchange in the C-terminal portion of the prodomain α2-helix in the TGF-β2 straitjacket (Fig. 2). In structures of latent TGF-β1 and GDF8, the α2-helix nestles against the convex side of the GF β6′-β7′ region (Fig. 4A,B). Although the corresponding peptide was not observed in HDX for TGF-β1, this region in latent GDF8 and all non-latent members is more rapidly deuterated suggesting a key difference in how the α2-helix is engaged specifically in TGF-β2. Secondly, a peptide (INPEASA) mapping to the α3′-β5′ loop exhibits slow exchange in TGF-β2 (Fig. 2). Although the corresponding peptide was not recovered in TGF-β1, in TGF-β1 structures the corresponding peptide forms varying H-bond networks with the sidechains of Arg-45 and Gln-52 in the α1-helix (Fig. 5C) and thus could interact with and stabilize the α1-helix. Furthermore, a Cys residue is present between the α2-helix and the fastener in TGF-β2 but not in TGF-β1 (arrow in Fig. 2). This Cys residue likely disulfide links to its partner in the other prodomain. The fastener forms important interactions with the α1-helix and completes the encirclement of the growth factor β6′-β7′ finger by the straitjacket. Disulfide linkage between the straitjackets in each monomer would cooperatively stabilize them.

**Figure 5.**
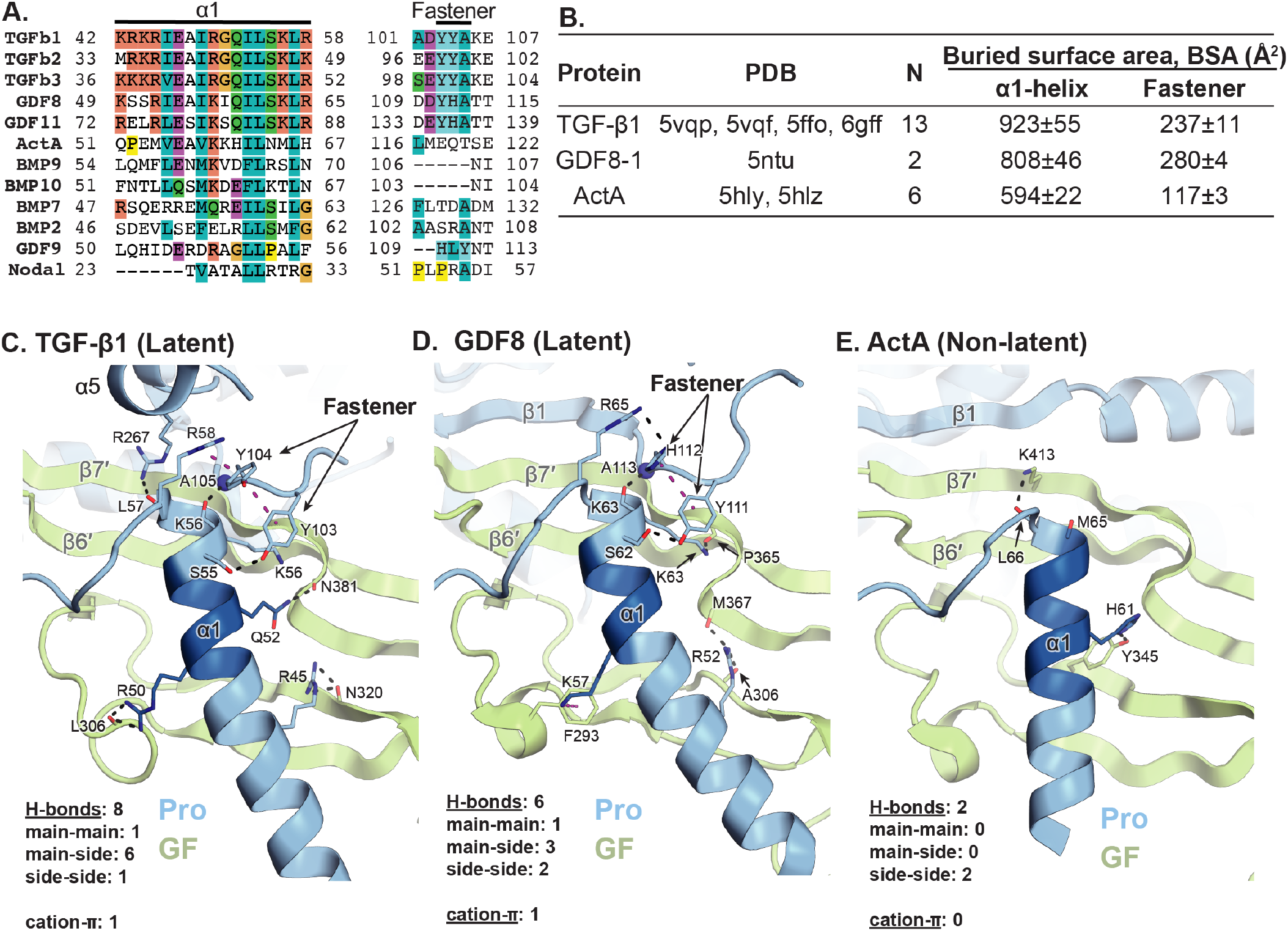
The α1-helix, fastener, and their interaction networks. **A.** Sequence alignment of α1-helix and fastener regions of representative family members. **B.** Solvent accessible surface area buried on the α1-helix and the fastener, using the overlined residues shown in panel A, as described in Results. For TGF-β1 and ActA, data are mean and s.d. of all independent monomeric units in the listed PDB accessions. For GDF8, data are mean and difference from the mean of each independent monomeric unit in the listed PDB accession. **C–E.** Ribbon cartoons of the α1-helix and its environment in TGF-β1 (*C*, PDB code 5vqp, chains A and B), GDF8 (*D*, PDB code 5ntu, chains A and B), and ActA (*E*, PDB code 5hlz, chains A, B, and C). Hydrogen and pi bonds to the α1-helix as well as pi bonds within the fastener are shown as black and magenta dashes, respectively. The backbone amide of A105 (C) and A113 (D) are shown as blue spheres.

We further examined crystal structures of TGF-β family members for structural correlates of slower exchange of the prodomain α1-helix in latency. Buried solvent accessible surface area correlates well with stability of interactions between proteins; therefore, we calculated burial for α1-helix and fastener residues (overlined in Fig. 5A). We measured burial of these residues by other residues in the prodomain and the growth factor monomer that is surrounded by the straitjacket. However, we omitted burial by the other growth factor monomer. This monomer was ordered in some, but not in other independent TGF-β1 and GDF8 monomers in the crystal asymmetric unit, showing that it does not strongly interact with the α1-helix. Buried solvent accessible surface area for the α1-helix and fastener was similar for TGF-β1 and GDF8 and substantially greater than for ActA (Fig. 5B). We also enumerated hydrogen bond and cation-pi interactions that the α1-helix made with the growth factor or prodomain (Fig. 5C–E). Latent TGF-β1 and GDF8 structures contained nine and seven such interactions, respectively, compared to two in ActA (Fig. 5C–E). A similar number of corresponding interactions were observed in other monomers in crystal asymmetric units and in other crystal structures of porcine and human TGF-β1, GDF8, and ActA.

All hydrogen bond and cation-pi interactions were found in the C-terminal portion of the α1-helix where HDX was low, from Arg-45 to Arg-58 in TGF-β1 and from Arg-52 to Arg-65 in GDF8 (Fig. 5A,C,D). The sidechains of homologous residues TGF-β1 Arg-45 and GDF8 Arg-52 form similar hydrogen bonds to their growth factor backbones (Fig. 5C,D). The sidechains of homologous residues TGF-β1 Arg-50 and GDF8 Lys-57 extended from the opposite face of the α1-helix to interact with another portion of the GF. Although they interacted with similar regions of the GF, they formed chemically distinct interactions. TGF-β1 Arg-50 formed bidentate hydrogen bonds to the GF backbone and GDF8 Lys-57 formed a cation-pi bond to GF residue Phe-293. Homologous α1-helix residues Ser-55 in TGF-β1 and Ser-62 in GDF8 form identical sidechain-sidechain hydrogen bonds to a Tyr residue in their fasteners.

“Caps” stabilize α-helices by forming hydrogen bonds to residues at their ends. Except at the ends, α-helical residues hydrogen bond through both their amide NH and CO groups to other α-helical residues. Residues at the C-terminal ends of helices form helical H-bonds through their NH but not CO groups. Hydrogen bonds donated by other structural elements to these unsatisfied helical residue carbonyl oxygens “cap” helices and stabilize them. TGF-β1 α1-helix residue Lys-56 is capped by a mainchain-mainchain hydrogen bond with fastener residue Ala-105, while α1-helix residue Leu-57 is capped by a mainchain-sidechain hydrogen bond with Arg-267 in the α5-helix (Fig. 5C). In GDF8, α1-helix residue Lys-63, which is homologous to Lys-56 in TGF-β1, is similarly capped by a mainchain-mainchain hydrogen bond with fastener residue Ala-113 (Fig. 5D). In contrast, homologous α1-helix residue Met-65 in ActA is left uncapped, with an unsatisfied CO group (Fig. 5E). This is because there is no structural equivalent to the fastener in ActA; instead, in a similar position in the sequence, an α-helix forms that locates 10 Å further away. Although the mainchain carbonyl oxygen at residue 66 of ActA forms a hydrogen bond, it hydrogen bonds to a Lys sidechain, not to backbone (Fig. 5E). In TGF-β1 and GDF8, other interactions stabilize the α1-helix interface with the fastener. In TGF-β1, Arg-58 forms a cation-pi bond with fastener residue Tyr-104 and in turn Tyr-104 and Tyr-103 form a T-shaped pi-pi interaction. In GDF8, homologous residues similarly interact. Arg-65 hydrogen bonds to His-112 and in turn His-112 forms a T-shaped pi-pi interaction with Tyr-111 (46). The interactions between the fastener and α1-helix in TGF-β1 and GDF8 secure the α1-helix to the remainder of the prodomain and complete straitjacket encirclement of the growth factor.

Our HDX studies demonstrate that slow exchange in the C-terminal end of the α1-helix correlates with latency and with the structural environment of this portion of the α1-helix in crystal structures. These results complement previous studies on the importance of the α1-helix in latency. Mapping studies of the minimum inhibitory prodomain fragment of GDF8 yielded fragments of varying lengths that all included the α1-helix (57–59). The shortest fragment was a 23-residue peptide that encompassed the entire α1-helix of GDF8 (59); however, the IC_50_ of this peptide (~10 μM) was 100-fold higher than that of the intact prodomain (1 nM) (30) indicating that other prodomain elements must also contribute. Notably, replacement with TGF-β1 fastener sequences in an ActA prodomain dimerized in an Fc fusion construct strongly inhibited the ActA growth factor (60). This result strongly supports our conclusion that fastener interactions with the α1-helix are important in conferring latency.

Mutational studies have amply illustrated the importance of the α1-helix and the fastener for latency. Mutation to alanine of hydrophobic residues Ile-53, Leu-54, Leu-57, and Leu-59 and basic residues Arg-45, Arg-50, Lys-56, and Arg-58 in the α1-helix all led to increased release of the TGF-β1 growth factor (61). Notably all four basic residues participate in interactions at the α1-helix–fastener–GF interface; Arg-45 and Arg-50 form polar interactions with the growth factor, whereas Arg-58 forms a cation-pi interaction with fastener (Fig. 5C). Mutation of TGF-β1 fastener residues Tyr-103 or Tyr-104 to Ala abolished latency (47). Furthermore, mutation of Lys-56 and Tyr-104 to Cys abolished expression or latency, respectively, while their double mutation restored expression and latency, suggesting formation of a disulfide bond. Moreover, the double mutation abolished activation by integrin αVβ6 but not by heat. In GDF8, alanine mutations of equivalent fastener residues Tyr-111 and His-112 and of the α1-helix residue Arg-65, which forms an H-bond with His-112 (Fig. 5D), increased basal activity over the wild type latent form (46,62).

Sequence alignments revealed remarkably strong sequence conservation in the α1-helix of all latent family members (e.g., TGF-β1–3, GDF8, and GDF11) with a consensus R(I/V/L)E(A/S)(R/K)I(R/K)XQILSKL(R/K) sequence (Fig. 5A). As discussed in the previous paragraph, the C-terminal portion of this sequence, and especially its hydrophobic and basic residues, have been shown to be important for latency in TGF-β1 and GDF8. In contrast, this region is much more divergent in non-latent family members, with only two hydrophobic residues in the consensus sequence shared with representative non-latent members (Fig. 5A) (55). Moreover, the fastener regions of latent members are all characterized by conserved, adjacent tyrosine and histidine aromatic residues that can participate in pi-pi interactions with one another and with α1-helix basic residues in cation-pi interactions (Fig. 5A). Strong sequence conservation of these signatures for the α1-helix and fastener suggest that similar interactions are present in latent TGF-β2, TGF-β3, and GDF11 and that prodomain association is sufficient to confer latency only for TGF-β1–3, GDF8, and GDF11.

In vivo, the bioavailability and activity of many family members (both latent and non-latent) are regulated by interactions with “milieu molecules” that include proteoglycans, ECM components, and cell surface transmembrane proteins (6–17). Our HDX studies highlight the association region at the N-terminus of the prodomain as a potential region to mediate these interactions. The interaction of TGF-βs with milieu molecules has been best characterized. A conserved Cys in the association region of TGF-βs forms disulfide linkages to latent TGF-β binding proteins in the ECM and the transmembrane proteins glycoprotein A repetitious predominant (GARP) and leucine rich repeat containing protein 33 (LRRC33) on cell surfaces. Rapid exchange of the association region in TGF-β1 and 2 suggest that in solution and in the absence of binding to a milieu molecule, the association region is exposed and unstructured. With the exception of BMP9, the prodomain segment N-terminal to the α1-helix in all family members studied here are in fast exchange, suggesting their availability for binding to as yet unknown factors. Of note, Cys residues are found in the putative association regions of GDF8 and ActA (Fig. 2) raising the possibility that they may form disulfide complexes with milieu molecules analogous to the TGF-βs.

Taken together, our EM and HDX-MS results point to rich diversity in the overall conformation, dynamics, and structural details (including prodomain–growth factor interfaces) in the TGF-β family. Moreover, structural dynamics investigation by HDX-MS has provided important context for interpreting existing crystal structures and insights into functional differences in the family. In particular, our HDX results coupled with strong sequence conservation of the α1-helix and fastener suggests that prodomain–GF association is sufficient for conferring latency only for TGF-β1–3 and GDF8 and 11, whereas latency in other family members may require association with a binding partner in the extracellular milieu.

## EXPERIMENTAL PROCEDURES

### Protein Expression and Purification

Human TGF-β2 was cloned into the pEF-puro vector with N-terminal 8xHIS and streptavidin-binding peptide (SBP) purification tags (47), a C24S mutation, an N140R mutation to remove one N-glycan site, and abolition of furin cleavage by replacing the residues 298–302 (RRKKR) with G. This TGF-β2 construct was stably expressed in 293S GnTI^−/−^ cells to produce protein with high-mannose glycosylation, purified by His and Streptactin affinity chromatographies, and dialyzed into 20mM Tris-HCl pH 7.5, 500 mM NaCl with Precision3C protease to remove purification tags. Cleaved TGF-β2 was then purified by Superdex200 size-exclusion chromatography (SEC) in 20mM Tris-HCl pH 7.5, 500 mM NaCl.

Full-length human ActivinA (wild type) and full-length BMP10 carrying N67Q and N131Q N-glycosylation site mutations and replacement of residues 312–316 (ARIRR) with glycine to abolish furin cleavage were cloned into the S2-2 vector (ExpreS2ion Biotechnologies) with N-terminal 8xHis and SBP tags and stably integrated into *Drosophila* S2 cells. Cells were adapted to growth in serum-free Excell 420 media. After 4 days, culture supernatant was collected, filtered, buffer exchanged to 20mM Tris-HCl pH 7.5, 500 mM NaCl and loaded onto a Ni-NTA column (Qiagen). The column was washed with 20 mM Tris, pH 7.5, 500 mM NaCl and 20 mM imidazole, and protein was eluted with 20 mM Tris-HCl, pH 7.5, 500 mM NaCl, and 1 M imidazole. Pooled elution fractions were dialyzed into 20 mM Tris, pH 7.5, 500 mM NaCl and simultaneously cleaved with Precision3C to remove the His-SBP tag. Cleaved ActA and BMP10 protein samples were then dialyzed against 20 mM Tris-HCl pH 7.5, 150 mM NaCl and subjected to another round of Ni-NTA chromatography to purify away uncleaved material. The flow-through was then loaded onto a Superdex 200 column equilibrated with 20 mM Tris-HCl pH 7.5, 150 mM NaCl for SEC.

### Negative stain electron microscopy

Purified TGF-β2 and BMP10 were subjected to SEC in 20 mM Tris-HCl pH 7.5, 150 mM NaCl immediately prior to negative-stain EM analysis to remove any aggregates. The peak fractions were loaded onto glow-discharged carbon-coated grids, buffer was wicked off, and grids were immediately stained with 0.75% (wt/vol) uranyl formate and imaged with an FEI Tecnai T12 microscope and Gatan 4K×4K CCD camera at 52,000× magnification (2.13 Å pixel size at specimen level) with a defocus of −1.5 μm. Well-separated particles (>5,000) were interactively picked using EMAN2 (63). Class averages were calculated by multireference alignment followed by K-means clustering using SPIDER (45,64–66).

### Hydrogen deuterium exchange mass spectrometry (HDX-MS)

HDX-MS studies were performed using methods reported previously (45,67). Additional experimental details are provided in Table S1 per the recommended format (68). ActA, BMP10, and TGF-β2 were expressed and purified in this study, whereas wild type human BMP7, a BMP9 chimera composed of the mouse prodomain and human growth factor domain, and human TGF-β1 carrying a C4S mutation, N-glycosylation site mutations N107Q and N147Q, and a R249A mutation to abolish furin cleavage were expressed and purified in other studies (23,48). All HDX-MS experiments were performed with proteins buffer exchanged into 20 mM Tris-HCl, pH 7.5, 150 mM NaCl with the exception of BMP9, which was buffer exchanged into PBS (1.8 mM KH_2_PO_4_, 10 mM Na_2_HPO_4_, 137 mM NaCl), pH 7.4. Samples (3 μL) of ActA (120.4 μM), BMP7 (272 μM), BMP9 (17 μM), BMP10 (60 μM), TGF-β1 (30 μM), and TGF-β2 (29 μM) were individually diluted 15-fold into 20 mM Tris, 150 mM NaCl, 99% D_2_O (pD 7.5) at room temperature for deuterium labeling. At time points ranging from 10 sec to 240 min, aliquots were removed and deuterium exchange was quenched by adjusting the pH to 2.5 with an equal volume of cold 150 mM potassium phosphate, 0.5 M tris (2-carboxyethyl) phosphine hydrochloride (TCEP-HCl), H_2_O. Each sample was analyzed as previously described (67,69). Briefly, the samples were digested online using a Poroszyme immobilized pepsin cartridge (2.1 mm x 30 mm, Applied Biosystems) at 15 °C for 30 s, then injected into a custom Waters nanoACQUITY UPLC HDX Manager™ and analyzed with a XEVO G2 mass spectrometer (Waters Corp., USA). The average amount of back-exchange using this experimental setup was 30–35%, based on analysis of highly deuterated peptide standards. All comparison experiments were done under identical experimental conditions such that deuterium levels were not corrected for back-exchange and are therefore reported as relative (70). The error of measuring the mass of each peptide averaged ± 0.15 Da in this experimental setup. All measurements were performed in technical duplicate.

Peptides were identified from triplicate high definition collision-induced dissociation mass spectrometry (HDMS^E^) analyses and data were analyzed using ProteinLynx Global SERVER (PLGS) 3.0.1 (Waters Corporation). A database containing only the sequences from human INHBA (ActA; Uniprot P08476), human BMP7 (Uniprot P18075), human BMP10 (Uniprot O95393), mouse GDF2 (BMP9; Uniprot Q9WV56) residues 23–318, human GDF2 (BMP9; Uniprot Q9UK05) residues 320–429, human GDF8 (Uniprot O14793), human TGFB1 (Uniprot P01137), and human TGFB2 (Uniprot P61812) was used with no cleavage specificity and no PTMs considered. Peptide masses were identified using a minimum of 130 ion counts for low energy threshold and a 50-ion count threshold for their fragment ions. The peptides identified in PLGS were then analyzed in DynamX 3.0 (Waters Corporation) implementing a minimum products per amino acid cut-off of 0.2, 2 consecutive product ions, and a maximum MH^+^ error of 10 ppm. Those peptides meeting the filtering criteria were further processed by DynamX to calculate relative % deuteration and display it on the sequence and tertiary structure.

## Data Availability

All HDX MS data have been deposited to the ProteomeXchange Consortium via the PRIDE (71) partner repository with the dataset identifier PXD026841.

## Supporting Information

This article contains supporting information (45).

## Acknowledgements

The authors would like to thank Melissa Chambers and Zongli Li for help with EM data collection, Jordan Anderson and Yang Su for critical discussions, and Margaret Nielsen for her assistance in designing figures. We acknowledge a research collaboration with the Waters Corporation (J.R.E.).

## Funding and additional information

This work was supported by NIH Grants R01-CA210920 and R01-AR67288 (T.A.S.). V.Q.L. was supported by NIH 5T32DK007527-35. Y.T. was supported by a Komen postdoctoral research fellowship (Komen PDF15334161).

## Conflict of Interest

The authors declare that they have no conflicts of interest with the contents of this article.

